# Lipopolysaccharide preconditioning augments phagocytosis of malaria-parasitized red blood cells through induced bone marrow-derived macrophages in the liver, thereby increasing the murine survival after *Plasmodium yoelii* infection

**DOI:** 10.1101/2020.05.22.111765

**Authors:** Takeshi Ono, Yoko Yamaguchi, Hiroyuki Nakashima, Masahiro Nakashima, Takuya Ishikiriyama, Shuhji Seki, Manabu Kinoshita

## Abstract

Malaria remains a grave concern for humans, as effective medical countermeasures for malaria infection have yet to be developed. Phagocytic clearance of parasitized red blood cells (pRBCs) by macrophages is an important front-line innate host defense against malaria infection. We previously showed that repeated injections of low-dose lipopolysaccharide (LPS) prior to bacterial infection, called LPS preconditioning, strongly augmented phagocytic/bactericidal activity in murine macrophages. However, how LPS preconditioning prevents murine malaria infection is unclear. We investigated the protective effects of LPS preconditioning against lethal murine malaria infection, focusing on CD11b^high^ F4/80^low^ liver macrophages, which are increased by LPS preconditioning. Mice were subjected to LPS preconditioning by intraperitoneal injections of low-dose LPS for 3 consecutive days, and 24 h later, they were intravenously infected with pRBCs of *Plasmodium yoelii* 17XL. LPS preconditioning markedly increased the murine survival and reduced parasitemia, while it did not reduce TNF secretions, only delaying the peak of plasma IFN-γ after malaria infection in mice. An *in vitro* phagocytic clearance assay of pRBCs showed that the CD11b^high^ F4/80^low^ liver macrophages of the LPS-preconditioned mice had significantly augmented phagocytic activity against pRBCs. The adoptive transfer of CD11b^high^ F4/80^low^ liver macrophages from LPS-preconditioned mice to control mice significantly improved the survival after malaria infection. We conclude that LPS preconditioning stimulated CD11b^high^ F4/80^low^ liver macrophages to augment the phagocytic clearance of pRBCs, which may play a central role in resistance against malaria infection. LPS preconditioning may be an effective tool for preventing malaria infection.

## Introduction

According to the ‘World Malaria Report 2019’ from the World Health Organization (WHO), malaria is one of the three major infectious diseases, along with acquired immune deficiency syndrome (AIDS) and tuberculosis, and efforts are underway to eliminate it (1). However, an increases in drug resistance to parasites and insecticide resistance to mosquitoes has threatened to worsen the infection control of malaria, leading to a high mortality (2, 3). The development of an effective malaria vaccine remains challenging (4). Therefore, there is an urgent need to establish effective medical countermeasures, including preventative and mitigation efforts, against severe malaria infection that do not rely on antimalaria drugs.

The augmentation of the host defense against malaria infections is important for improving patient outcomes. If activation of the innate immunity is effective for eliminating malaria parasites, a reduced malaria mortality can be expected, and instances of drug-resistant malaria can be eliminated. However, previously reported methods of activating innate immunity may also induce an enhanced inflammatory response in the hosts, resulting in organ damage (5). Augmentation of the elimination of infected cells without enhancing the inflammatory response may be an ideal medical countermeasure against malaria infection.

We recently reported that repeated low-dose LPS injection, termed LPS preconditioning or LPS tolerance (6) (7), renders mice drastically resistant to bacterial infection, as such LPS preconditioning potently reduces the host’s inflammatory response to bacterial stimuli while markedly augmenting the host’s bactericidal activity (8). LPS preconditioning increases the population of monocyte-derived macrophages in the liver and enhances their phagocytosis and bactericidal activity.

The liver plays a crucial role in malaria infection in the initial phase. We therefore attempted to use these attractive phenomena induced by LPS preconditioning as novel medical countermeasures against malaria infection and investigated whether or not LPS preconditioning can prevent severe malaria infection using a murine malaria infection model.

## Results

### LPS preconditioning improved the mouse survival after lethal malaria infection

LPS preconditioning was induced by intraperitoneal injections of 5, 50 or 500 μg/kg of LPS for 3 consecutive days in mice, and 24 h after the last LPS injection, they were intravenously infected with the GFP-expressing *Plasmodium yoelii* 17XL (PyLGFP) (5 × 10^4^ pRBCs). This dose of PyLGFP infection was lethal for non-treated control mice (Fig. 1A-C). However, preconditioning with 5 μg/kg of LPS tended to prolong the murine survival time after PyLGFP infection (Fig. 1A). Interestingly, preconditioning with 50 μg/kg of LPS showed a 20% survival rate in PyLGFP-infected mice (Fig. 1B), and preconditioning with 500 μg/kg LPS resulted in more than half of the infected mice surviving (60% survival, Fig. 1C), suggesting that LPS preconditioning significantly increased the survival of PyLGFP-infected mice.

**FIG 1.**
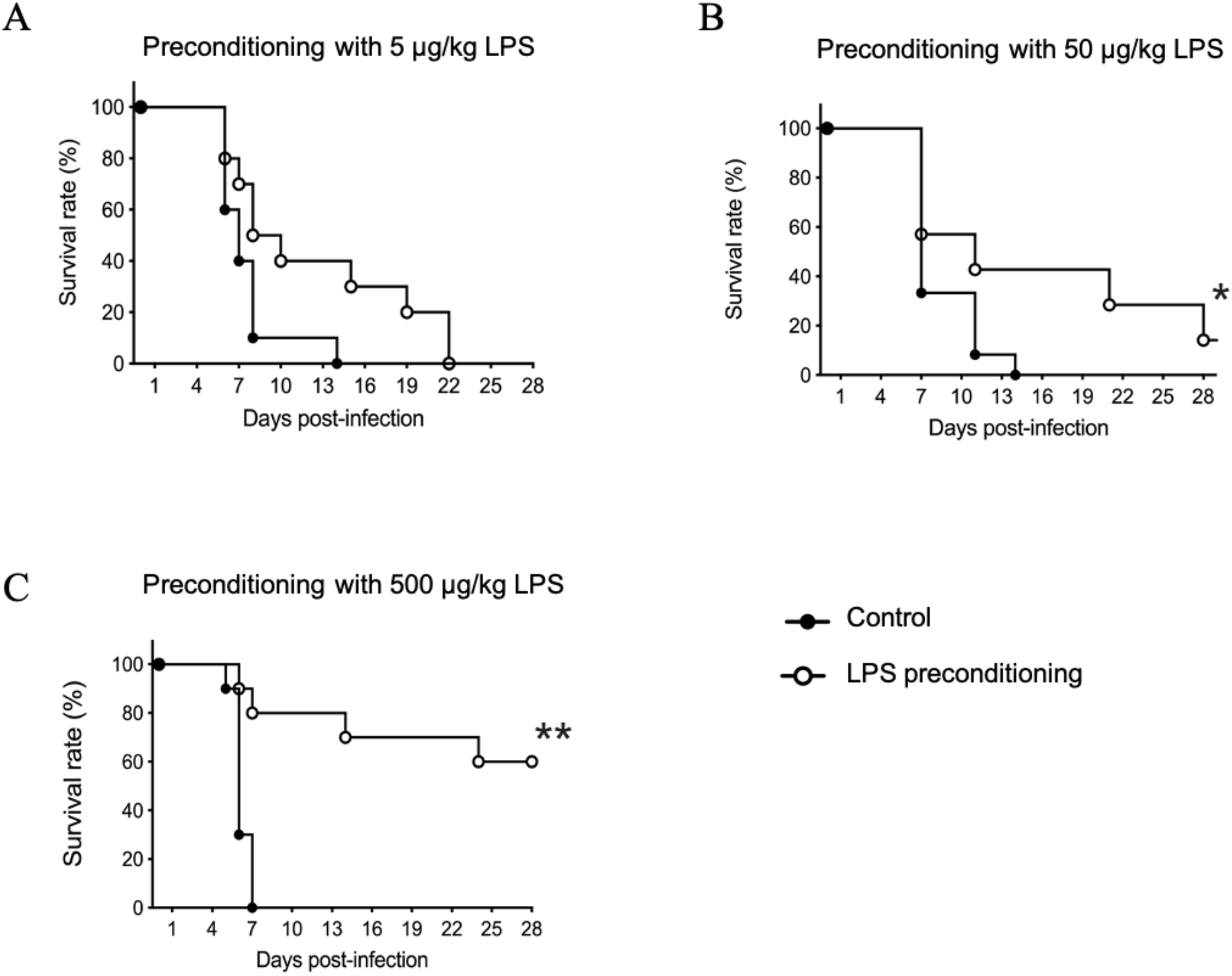
The survival after rodent malaria infection in LPS-preconditioned mice. LPS preconditioning was induced in mice with i.p. injection of 5 (A), 50 (B), or 500 (C) μg/kg LPS for 3 consecutive days, and 24 h after the last LPS injection, the mice were i.v. infected with 5 × 10^4^ pRBCs of PyLGFP. Ten mice in each group. **p < 0.01, *p < 0.05 vs. control.

### LPS preconditioning reduced the growth of malaria in mice at 5 days after infection

Next, we examined the effect of LPS preconditioning (5 or 500 μg/kg of LPS) on the growth of PyLGFP in mice. Although no significant reduction was noted at 3 days, LPS preconditioning with both 5 and 500 μg/kg LPS significantly reduced the parasitemia at 5 days after infection (Fig. 2A, B). We confirmed the significant reduction in parasitemia by LPS preconditioning using a flow cytometric analysis. LPS preconditioning with both 5 and 500 μg/kg LPS significantly reduced the proportion of GFP-positive RBCs, which indicated PyLGFP-parasitized RBCs, at 5 days after infection (Fig. 2C, D). This suggested that LPS preconditioning potently reduced growth of malaria in mice during the early phase of parasite infection (at 5 days), resulting in an improved survival after infection.

**FIG 2.**
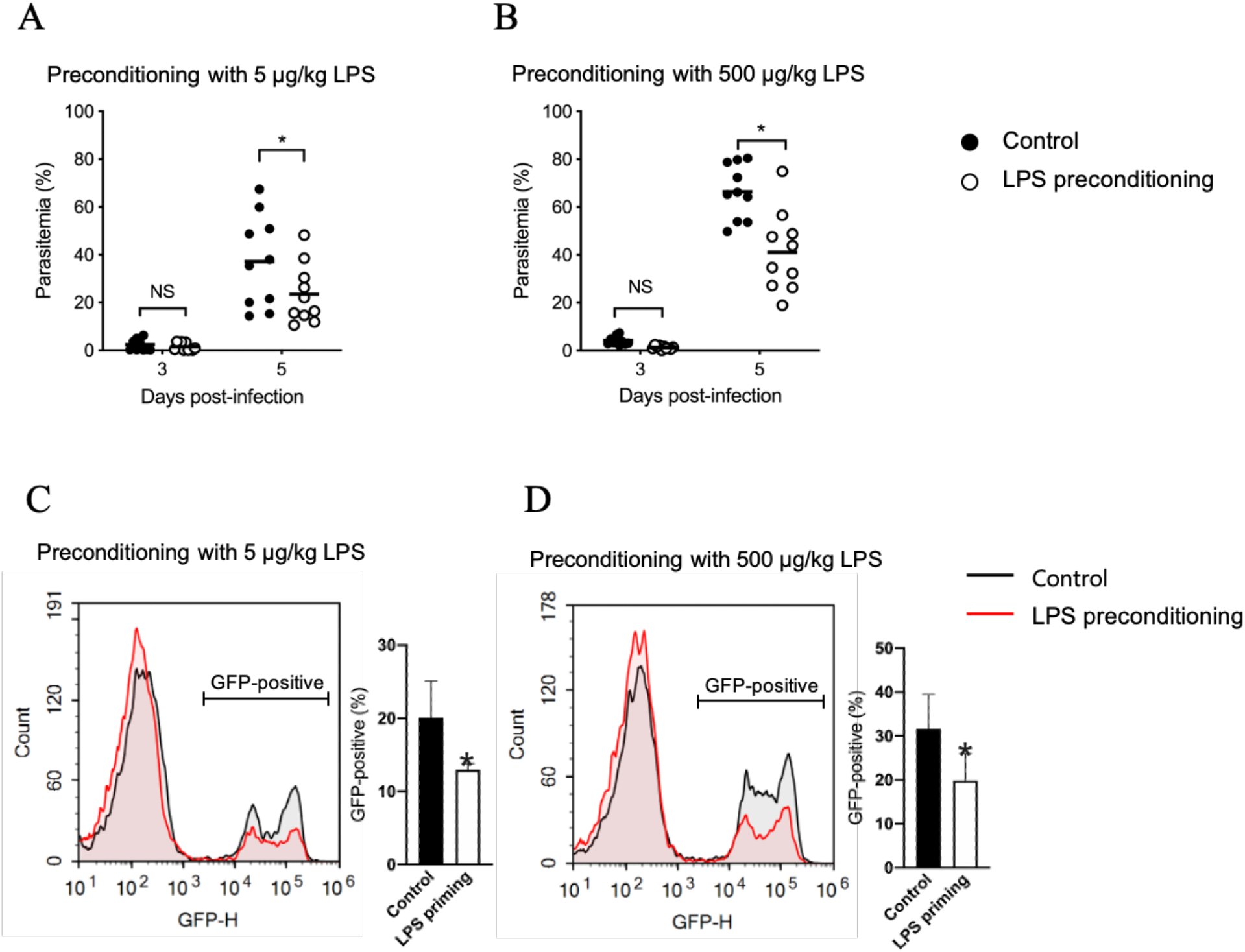
The effects of LPS preconditioning on the parasitemia after *Py*LGFP infection in mice. Mice were similarly subjected to LPS preconditioning with i.p. injection of 5 μg/kg or 500 μg/kg LPS, and 24 h later, they were i.v. infected with PyLGFP-parasitized RBCs. Parasitemia at days 3 and 5 post-infection was measured by Giemsa stain (A, B). The proportion of PyLGFP-parasitized RBCs was also evaluated at day 5 as the FITC intensity using a flow cytometer (C, D). Data are the means ± SE from 10 mice in each group. *p < 0.05 vs. control.

### LPS preconditioning augmented the phagocytic clearance of pRBCs by CD11b^high^ F4/80^low^ macrophages in the murine liver

We first examined the effect of LPS preconditioning on the phagocytic clearance of pRBCs in the murine liver mononuclear cells (MNCs). Liver MNCs were obtained from the LPS-preconditioned mice 24 h after the last LPS injection to use for the following examinations. After co-culture with pRBCs for 16 h, the liver MNCs of the LPS-preconditioned mice (with 500 μg/kg LPS) significantly reduced the number of pRBCs in the culture medium compared with the control mice (Fig. 3A), suggesting that LPS preconditioning augmented the phagocytic clearance of pRBCs by liver MNCs.

**FIG 3.**
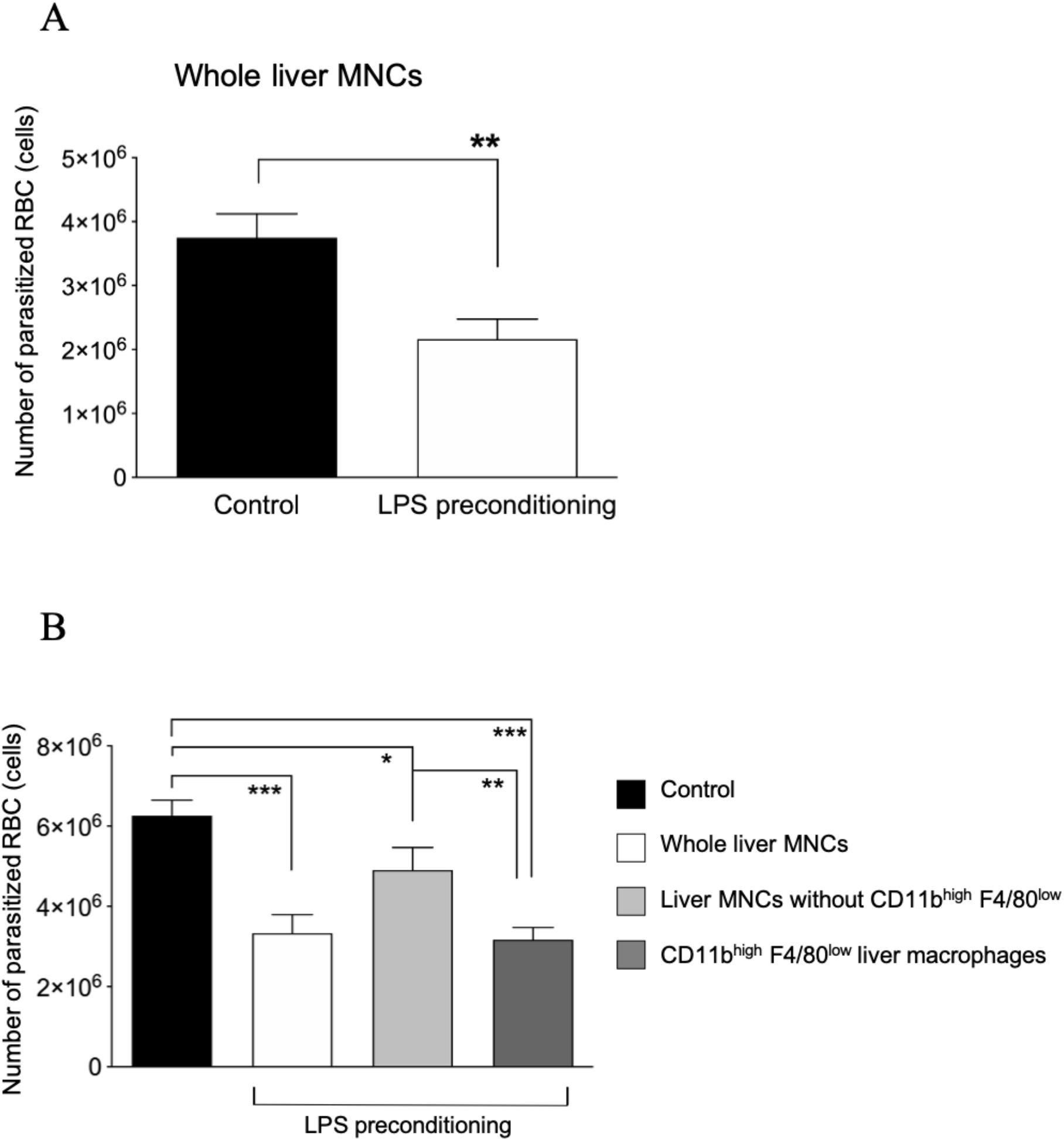
An *in vitro* phagocytosis assay. Liver MNCs (A) or sorted CD11b^high^ F4/80^low^ liver macrophages (B) were obtained from the LPS (500 μg/kg)-preconditioned mice and cocultured with PyLGFP-parasitized RBCs for 16 h. Thereafter, residual pRBCs were counted. Data are the means ± SE from 5 mice in each group. ***p < 0.0001, **p < 0.01, *p < 0.05.

Next, we examined the phagocytic clearance of pRBCs by the CD11b^high^ F4/80^low^ liver macrophages, which are markedly increased by LPS preconditioning (8). We sorted the CD11b^high^ F4/80^low^ liver macrophages and other liver MNCs of the LPS-preconditioned mice (with 500 μg/kg LPS) and co-cultured these cells (5 × 10^5^ cells) with pRBCs for 16 h. The CD11b^high^ F4/80^low^ liver macrophages of the LPS-preconditioned mice significantly reduced the number of residual pRBCs in the culture medium compared with other MNC subsets in the LPS-preconditioned mice (Fig. 3B), which subset however showed a significant reduction in residual pRBCs compared to the whole liver MNCs of the control mice [Sorry, this is unclear: please clarify the meaning of the highlighted text] (Fig. 3B). This suggested that LPS preconditioning stimulated liver MNCs, particularly CD11b^high^ F4/80^low^ liver macrophages, encouraging them to augment the phagocytic clearance of pRBCs.

### Changes in the plasma cytokine levels after malaria infection in mice

LPS preconditioning markedly reduces the TNF and IFN-γ secretion in mice after bacterial infection (8). We then examined these cytokine responses to malaria infection in the LPS-preconditioned mice. Unlike bacterial infection, no significant difference in the plasma TNF levels were observed after PyLGFP infection between the control and LPS-preconditioned mice with 5 or 500 μg/kg LPS (Fig. 4A, B). The plasma IFN-γ levels peaked at 3 days after PyLGFP infection in control mice, whereas LPS preconditioning with 5 and 500 μg/kg LPS both delayed those IFN-γ peaks, which instead peaked at 5 days after infection (Fig. 4C, D).

**FIG 4.**
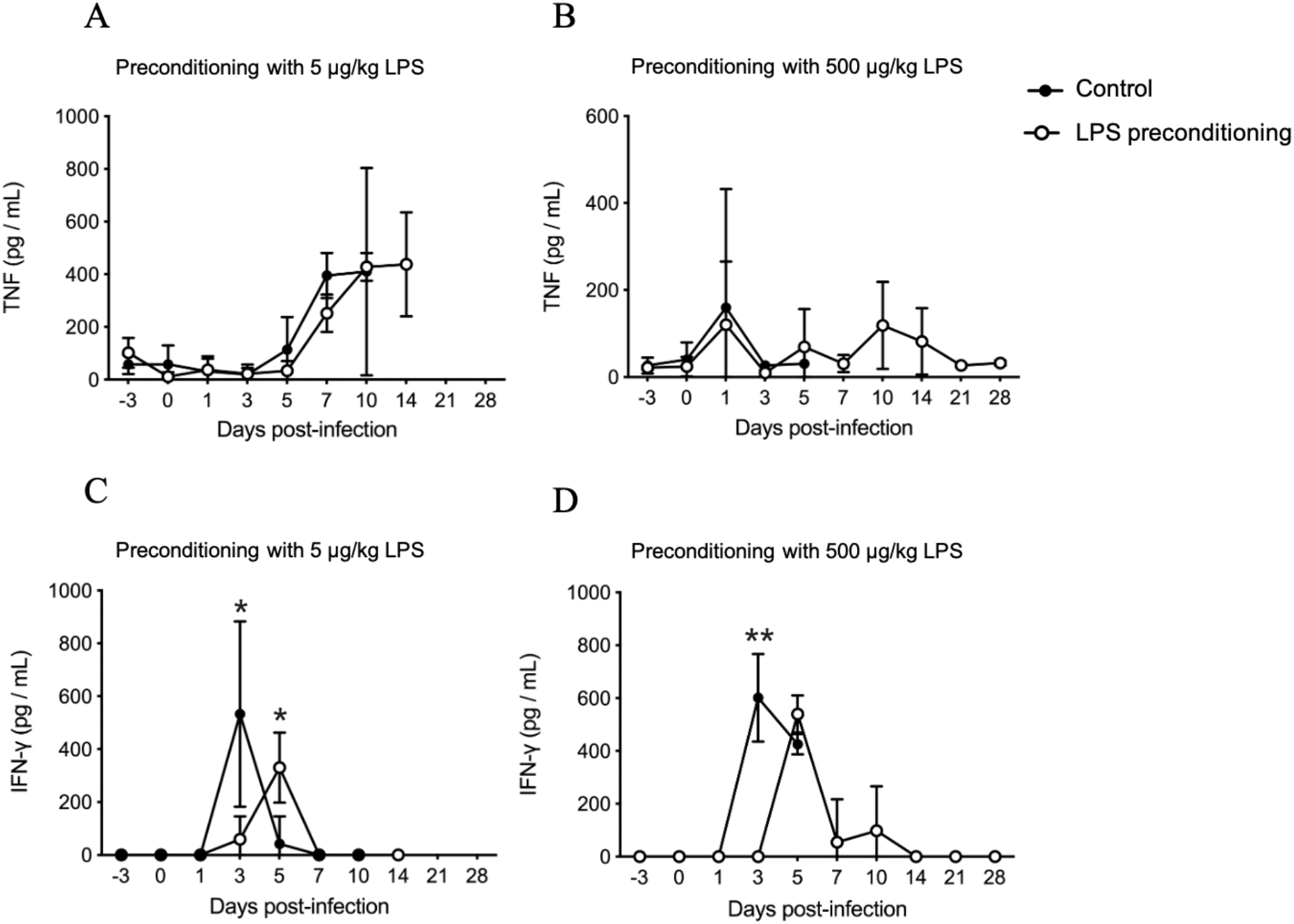
Changes in the plasma cytokine levels after PyLGFP infection in LPS-preconditioned mice and control mice. LPS-preconditioned mice with 5 μg/kg LPS (A, C) or 500 μg/kg LPS (B, D) and control mice were i.v. infected with 5 × 10^4^ PyLGFP-parasitized RBCs to examine the plasma IFN-γ and TNF. Data are the means ± SE from 10 mice in each group. **p < 0.01, *p < 0.05 vs. control (at the same time point).

### Pathological observation of the murine liver 5 days after malaria infection

In the pathological examination of hematoxylin-eosin (H.E.) staining, RBC-phagocytosis by macrophages was markedly observed in the liver of LPS-preconditioned mice (with 500 μg/kg LPS) compared with control mice (Fig. 5). To confirm the phagocytosis of PyLGFP-parasitized RBCs in the murine liver, we performed the immunohistochemical staining of GFP in the liver of PyLGFP-infected mice. Consistent with H.E. staining, positive staining of GFP in liver macrophages, which indicates phagocytosis of pRBCs, was also obviously observed in the LPS-preconditioned mice compared with the control mice (Fig. 6).

**FIG 5.**
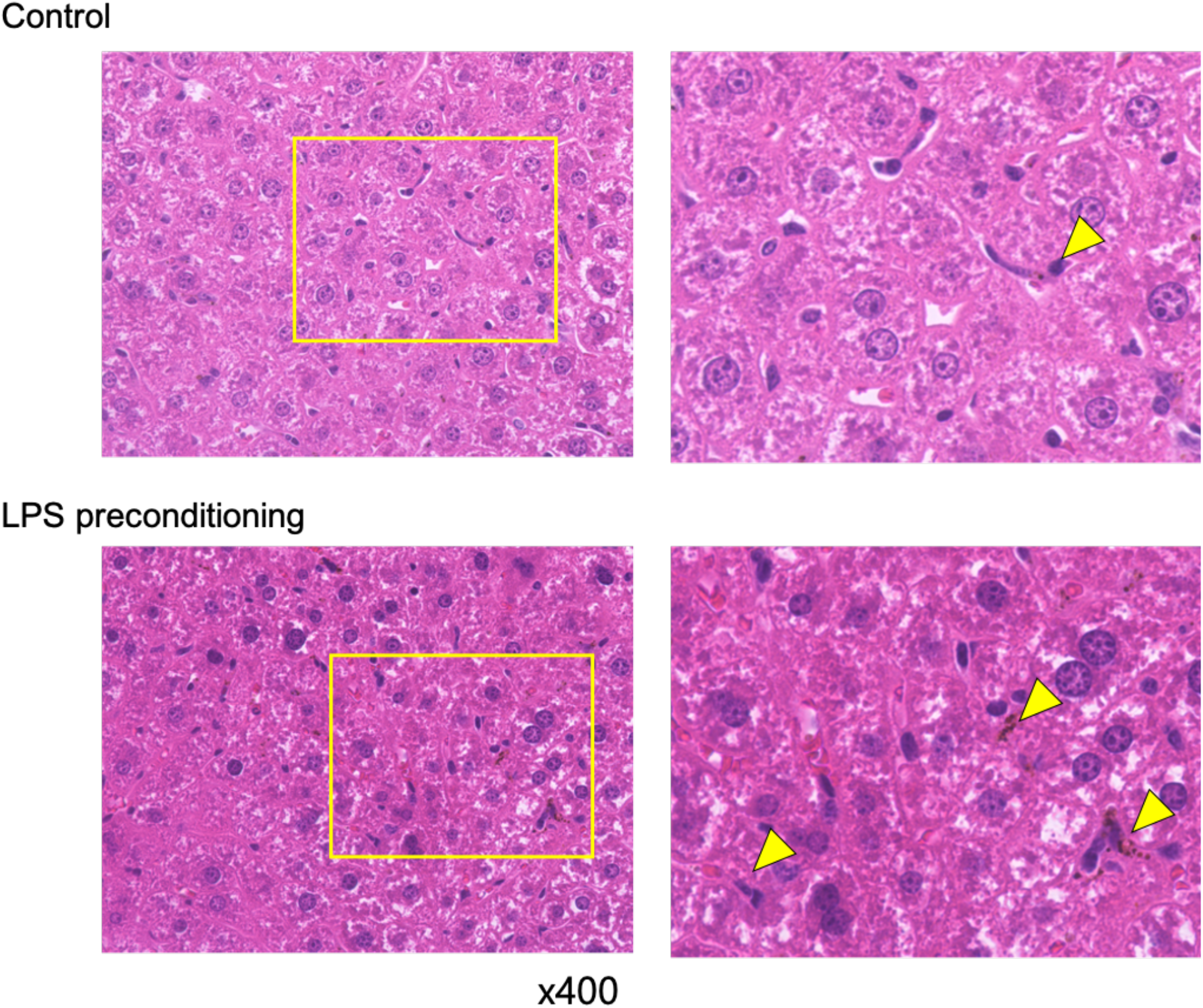
Results of a pathological examination of the liver after PyLGFP infection. Liver sections of LPS (500 μg/kg)-preconditioned mice and control mice were stained with H.E. Data are representative of three mice in each group with similar results. Right columns are magnified images of the yellow squares in the left columns. Arrowheads indicate RBC-phagocytosed macrophages.

**FIG 6.**
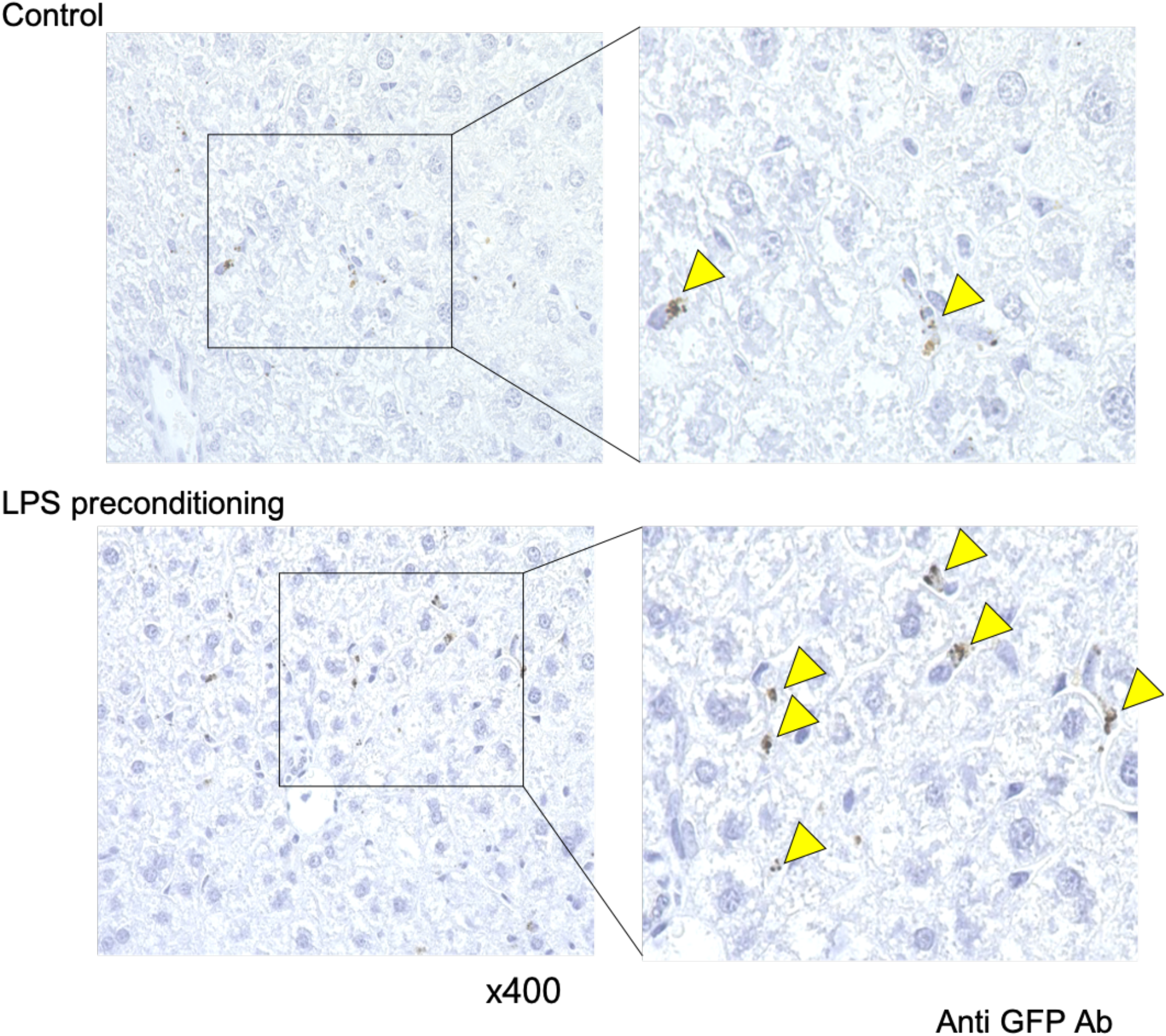
Results of an immunohistochemical analysis of pRBC-phagocytosed macrophages in the liver. Liver sections of LPS (500 μg/kg)-preconditioned mice and control mice were stained with anti-GFP Ab, as described in the Materials and methods. Arrowheads indicate cells with GFP-positive staining, which are pRBCs. Data are representative of three mice in each group with similar results. Right columns are magnified images of the squares in the left columns.

### Adoptive transfer of CD11b^high^ F4/80^low^ liver macrophages from LPS-preconditioned mice induced resistance to malaria infection in control mice

To confirm the effect of CD11b^high^ F4/80^low^ liver macrophages, which are induced by LPS preconditioning, on the resistance to rodent malaria infection, we performed the adoptive transfer of CD11b^high^ F4/80^low^ liver macrophages from the LPS-preconditioned mice (with 500 μg/kg LPS) to control mice and then infected these mice with PyLGFP. Adoptive transfer of CD11b^high^ F4/80^low^ liver macrophages from the LPS-preconditioned mice to the control mice significantly prolonged the survival time after malaria infection, although the adoptive transfer of other liver MNC subset cells did not improve the survival (Fig. 7A). However, there were no significant changes in plasma IFN-γ or TNF levels after PyLGFP infection among the three mouse groups (Fig. 7B, C). These results support the notion that CD11b^high^ F4/80^low^ liver macrophages, which are a key fraction in the induction of LPS preconditioning, play a major role in resistance to malaria infection.

**FIG 7.**
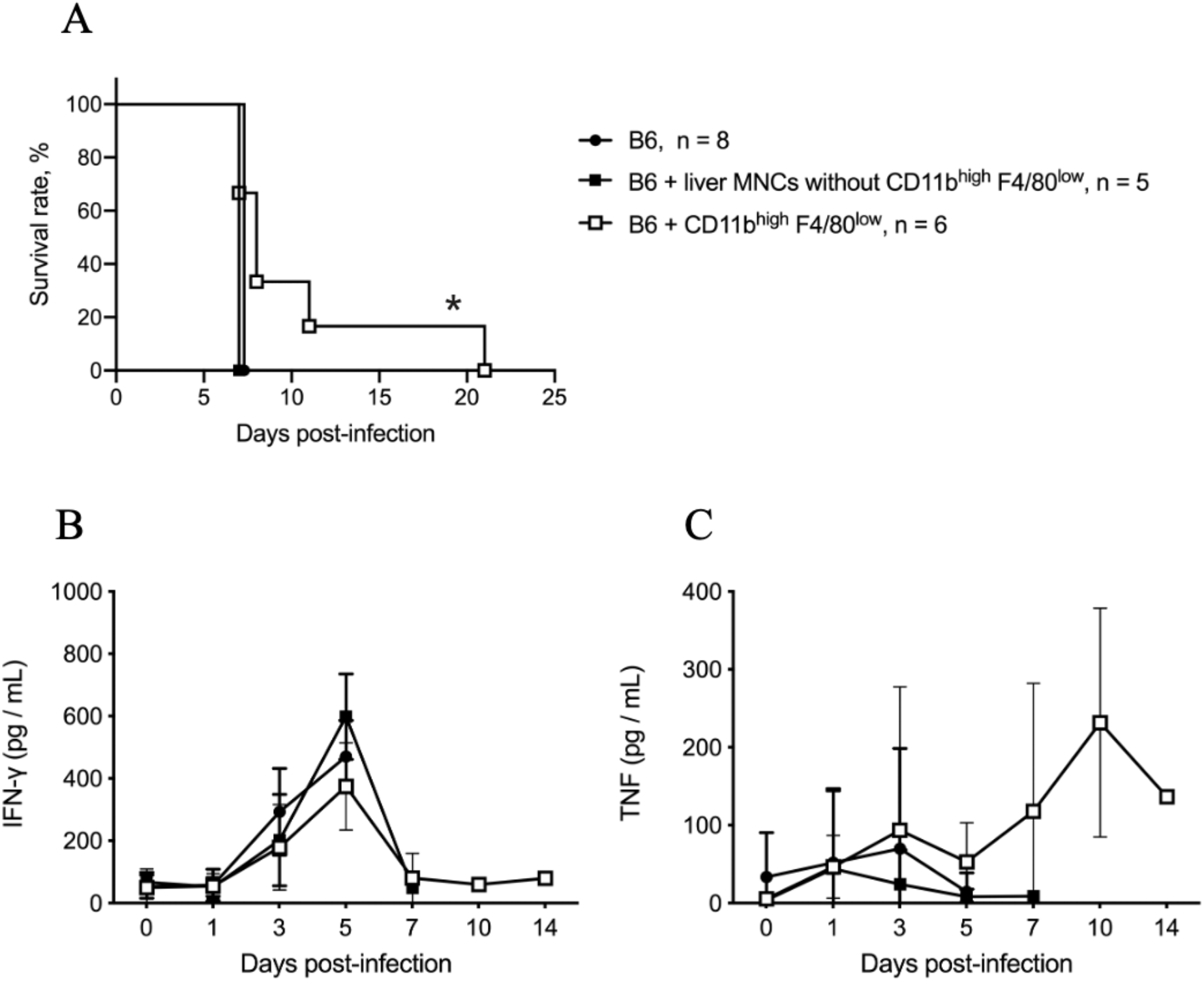
Adoptive transfer of CD11b^high^ F4/80^low^ liver macrophages from LPS-preconditioned mice to control mice. CD11b^high^ F4/80^low^ liver macrophages and CD45^+^ liver MNCs (except for the CD11b^high^ F4/80^low^ subset) were obtained from the LPS-preconditioned mice (with 500 μg/kg LPS) and transferred to control mice. Thereafter, mice were i.v. infected with PyLGFP-parasitized RBCs to monitor their survival (A), plasma IFN-γ (B), and TNF (C) levels. The number of mice in each group is indicated in the panel. * p < 0.01 vs. control mice and p < 0.05 vs. mice with transferred CD45^+^ liver MNCs except for CD11b^high^ F4/80^low^ cells.

## Discussion

LPS preconditioning significantly augmented the phagocytic clearance of PyLGFP-parasitized RBCs by CD11b^high^ F4/80^low^ liver macrophages (Fig. 3B), which are monocyte-derived macrophages increased by LPS preconditioning (8), resulting in the improvement of the murine survival after PyLGFP infection (Fig. 1). We confirmed the beneficial effect of CD11b^high^ F4/80^low^ liver macrophages on the survival of PyLGFP-infected mice through the adoptive transfer of these macrophages (Fig. 7). LPS preconditioning effectively prevented rodent malaria infection without enhancing the proinflammatory cytokine responses to malaria infection (Fig. 4). This approach may be an attractive medical countermeasure for malaria infection.

Although the T cell-mediated immune response to initial malaria infection is well described, other immune cells, such as dendritic cells and monocytes/macrophages, have been shown to modulate immune activation and the severity of disease as well (9, 10). However, mice lacking tissue-resident macrophages experience increased malaria-related complications, such as disruptions in the blood-brain barrier, increased vascular permeability in the liver, and increased accumulation of hemozoin pigment in the lung (11). These studies imply a critical role for macrophages in the initial response to malaria infection. The liver stage of malaria is the first phase of infection in the hosts, and the liver-resident macrophages, namely Kupffer cells, play important roles in ameliorating the severity of malaria infection and preventing parasite release into the blood circulation (12).

LPS preconditioning may stimulate the CD11b^high^ F4/80^low^ liver macrophages to augment phagocytic activity against bacteria, which are external pathogens (8). These monocyte-derived macrophages alter their phagocytic phenotype, like Kupffer cells, as LPS preconditioning strongly upregulates their Fc-γ receptor I (FcγRI) expression (8), which is a critical FcγR involved in phagocytosis (13). FcγRs are also closely involved in opsonization, which is the process of antibodies binding to the pathogen and enabling ingestion and elimination by macrophages. However, unlike common bacteria (*Escherichia coli, Pseudomonas aeruginosa*), infected malaria parasitizes host RBCs. To eliminate malaria infection, macrophages must effectively phagocyte malaria-parasitized RBCs. Interestingly, a recent study reported that splenic red pulp macrophages, which are distinct from monocyte-derived macrophages, effectively phagocytose IgG-opsonized RBCs using FcγRs, most notably FcγRI (13). The CD11b^high^ F4/80^low^ liver macrophages induced by LPS preconditioning, which may show an increased FcγRI expression on their surface albeit monocyte-derived ones [Sorry, this is unclear: please clarify the meaning of the highlighted text], may demonstrate enhanced phagocytosis of malaria-parasitized RBCs through upregulated FcγRI.

LPS preconditioning drastically reduces proinflammatory cytokine responses, such as TNF and IFN-γ, to bacterial stimuli (7, 8). However, LPS preconditioning did not decrease TNF secretion after PyLGFP infection in mice (Fig. 4). Macrophages do not robustly produce proinflammatory cytokines, including TNF, during the early phase of malaria infection because phagosomal acidification of malaria-parasitized RBCs prevents macrophages from producing proinflammatory cytokines (14). Nevertheless, malaria-parasitized RBCs opsonized by immune serum increase TNF secretion from macrophages, suggesting that TNF production by macrophages is closely involved in their FcγR-mediated phagocytosis of malaria (15). Unlike bacterial infection, which involves an external pathogen, macrophages do not produce large amounts of TNF due to malaria infection; macrophage-produced TNF may be necessary for effective phagocytic elimination of parasitized RBCs. In the current study, sustained (albeit a low amounts of) TNF secretion may have helped enhance the elimination of malaria-parasitized RBCs by macrophages induced by LPS preconditioning, resulting in an improvement in murine malaria infection.

IFN-γ is the most powerful proinflammatory cytokine. An IFN-γ-induced tissue inflammatory reaction is required for effective bacterial elimination by the host; however, an IFN-γ-induced exaggerated inflammatory response can also cause sepsis and multi-organ injuries (16). LPS preconditioning did not reduce the plasma peak of IFN-γ but did delay its peak from 3 days to 5 days after PyLGFP infection (Fig. 4). In murine malaria infection, neutralizing IFN-γ prolonged lethal rodent malaria infection but did not reduce parasitemia (17). Similar to sepsis, the presence of proinflammatory cytokines, including IFN-γ, during acute malaria infection may be related to signs of severe malaria pathogenesis (18), IFN-γ secretion may not (at least negatively) affect parasitemia in mice. However, the rationale of this delayed peak of IFN-γ induced by LPS preconditioning should be investigated further in a future study.

In conclusion, LPS preconditioning rendered mice resistant to rodent malaria infection. CD11b^high^ F4/80^low^ liver macrophages that are induced by LPS preconditioning enhance the phagocytic clearance of parasitized RBCs, thereby improving the survival in malaria-infected mice.

## Materials and Methods

The Ethics Committee of Animal Care and Experimentation in National Defense Medical College Japan approved all requests for animals and the intended procedures of the present study (Permission number:18025).

### Animals and reagents

Male C57BL/6 mice (8 weeks old, body weight 20 g) were purchased from Japan SLC (Hamamatsu, Japan) and used for this study. LPS (*E. coli* 0111:B4) was purchased from Sigma-Aldrich (St. Louis, MO, USA) to use for LPS preconditioning. The *P. yoelii* 17XL lethal strain has been kept in our laboratory as described elsewhere (19). We created a stable GFP-expressing *P. yoelii* 17XL (PyLGFP) by transfection of an uncloned population of *P. yoelii* 17XL parasites with the vector pL0016, as described previously (20, 21).

We obtained pRBCs of PyLGFP from donor mice after intravenous (i.v.) inoculation with a frozen stock of parasites. After i.v. injection of PyLGFP, we checked the parasitemia of the donor mice daily. Thereafter, the pRBCs of PyLGFP were obtained at the proliferation phase of parasitemia for use in the current experiment (parasite rates of PyLGFP were 10%-15% of RBCs).

### Infection of rodent malaria (PyLGFP) in mice

After obtaining pRBCs of PyLGFP from the donor mice, 2.5 × 10^5^ pRBCs/mL with 0.2 mL RPMI1640 medium were i.v. injected into recipient mice 24 h after the last injection of LPS (LPS-preconditioned mice). The percentage of pRBCs in the recipient mice was monitored by thin tail blood smears stained with Giemsa stain. The murine survival was monitored every day.

### *Induction of* in vivo *LPS preconditioning*

LPS preconditioning was induced in mice by an intraperitoneal (i.p.) injection of 5, 50, or 500 μg/kg of LPS (dissolved in 0.5 mL saline) once daily for three times, as we previously described (8). Control mice were similarly i.p. injected with saline (0.5 mL) three times.

### Analyses of PyLGFP-parasitized RBCs using flow cytometry

RBCs were obtained from the PyLGFP-infected mice at five days after infection. Percentages of GFP-positive RBCs were evaluated as FITC intensity using flow cytometer (ACEA Biosciences, San Diego, CA, USA). After gating RBCs, GFP-positive RBCs were detected as FITC-positive.

### Isolation of liver MNCs from mice

Liver MNCs, including macrophages, were obtained from mice as described elsewhere (22). In brief, under deep isoflurane anesthesia, the liver was removed and minced with scissors. After shaking with 10 mL of Hank’s balanced salt solution containing 0.05% collagenase (Type IV; Sigma-Aldrich) for 20 min at 37 °C, liver specimens were filtered through mesh, suspended in 33% Percoll solution (Sigma-Aldrich) containing 10 U/mL heparin, and centrifuged for 15 min at 500 × g at room temperature. After lysing RBCs, the remaining cells were washed twice to obtain the liver MNCs.

### Sorting CD11b^high^ F4/80^low^ liver macrophages

LPS preconditioning increases the number of CD11b^high^ F4/80^low^ macrophages in the murine liver and potently augments their bactericidal activity (8). To sort this CD11b^high^ F4/80^low^ subset from the liver MNCs, obtained liver MNCs were stained with FITC-conjugated anti-F4/80 monoclonal antibody (mAb) (clone BM8; eBioscience, San Diego, CA, USA), PE-conjugated anti-CD11b mAb (clone M1/70, eBioscience), and APC-conjugated anti-CD45 mAb (clone 30-F11, eBioscience). Thereafter, CD11b^high^ F4/80^low^ CD45^+^ cells were sorted using a Sony SH800 cell sorter (Sony, Tokyo, Japan) (Supplemental Fig. 1). We also sorted CD45^+^ MNCs except for the CD11b^high^ F4/80^low^ cell subset from the liver MNCs using a cell sorter.

### In vitro *phagocytic clearance of PyLGFP-parasitized RBCs by liver MNCs or CD11b^high^ F4/80^low^ liver macrophages*

To examine the pRBC phagocytic clearance by liver MNCs, liver MNCs (5 × 10^5^ cells/200 μL) were obtained from LPS-preconditioned mice 24 h after the last LPS injection or control non-treated mice and then cocultured with 1 × 10^7^ pRBCs of PyLGFP (PyLGFP parasite rates were 10%-15% of RBCs) in antibiotic-free RPMI1640 medium for 16 h. The CD11b^high^ F4/80^low^ liver macrophages or other CD45^+^ liver MNCs were sorted from the liver MNCs in mice 24 h after the last LPS injection. Thereafter, these sorted cells (5 × 10^5^ cells/200 μL) were similarly cocultured with 1 × 10^7^ pRBCs for 16 h. After coculture for 16 h, the total RBC count in the culture medium was measured, and the proportion of pRBCs was counted in the smear stained with Giemsa stain. We then obtained the number of residual PyLGFP-parasitized RBCs in the culture medium.

### Adoptive transfer of CD11b^high^ F4/80^low^ liver macrophages from the LPS-preconditioned mice to control mice

To examine the effect of CD11b^high^ F4/80^low^ liver macrophages that are induced by LPS preconditioning on the rodent malaria infection, CD11b^high^ F4/80^low^ CD45^+^ cells were sorted from the liver MNCs of the LPS-preconditioned mice using the cell sorter. The CD45^+^ liver MNCs except CD11b^high^ F4/80^low^ subset were also sorted from the liver MNCs of the LPS-preconditioned mice. Thereafter, these sorted cells (1 × 10^6^ cells /200 μL PBS) were adoptively transferred into the recipient normal mice, and 2.5 × 10^5^ of pRBCs (with 0.2 mL RPMI1640 medium) were subsequently i.v. injected into recipient mice.

### Measurements of plasma cytokines in mice

Blood samples were obtained from the mice via submandibular bleeding using 5-mm GoldenRod Animal Lancets (MEDIpoint Inc., Mineola, NY, USA). Plasma TNF and IFN-γ levels were measured using their respective ELISA kits (BD OptEIA™; BD Biosciences, San Diego, CA, USA).

### Pathological analyses of the liver in the rodent malaria-infected mice

Under deep isoflurane anesthesia, the livers were removed from the mice 5 days after PyLGFP infection. Isolated liver samples were fixed in 4% formaldehyde, embedded in paraffin and sectioned at a thickness of 3 μm. The sections were subjected to H.E. staining or an immunohistochemical analysis. Regarding immunohistochemical staining of GFP, anti-GFP polyclonal antibody (1:500; MBL, Nagoya, Japan) was used as the primary antibody, HRP-labeled anti-rabbit IgG antibody (Nichirei, Tokyo, Japan) was used as the secondary antibody, and diaminobenzidine was used for the colorimetric reaction. These procedures were outsourced to LSI Medience (Tokyo, Japan).

### Statistical analyses

Statistical analyses were performed using the GraphPad Prism software program, version 8 (GraphPad Software, San Diego, CA, USA). A two-way analysis of variance (ANOVA) followed by Bonferroni’s post-hoc comparison test was used to assess the percentage of parasitemia. A repeated-measure ANOVA was used to compare the plasma cytokine levels between control and LPS-preconditioned mice. The data were the means ± standard error (SE) and were analyzed using either Student’s *t*-test or Mann–Whitney U test. p < 0.05 was considered statistically significant.

## Acknowledgments

This work was supported by JSPS KAKENHI Grant Number 19K07493 (T.O. and M.K.).

